# Genomic Characterization of Tissue Invasive *Klebsiella pneumoniae* Complex in a Hospital System with a Focus on Species Distribution and Hypervirulence

**DOI:** 10.1101/2020.11.04.369173

**Authors:** Jessica Bohrhunter, Samantha Taffner, Jun Wang, Dwight Hardy, Nicole Pecora

## Abstract

*Klebsiella pneumoniae* is an opportunistic pathogen known for multidrug resistance. Current research indicates that *K. pneumoniae* is a complex of closely related organisms *(K. pneumoniae sensu stricto, K. quasipneumoniae, K. variicola, K. quasivariicola,* and *K. africana).* Additionally, some strains of *K. pneumoniae sensu stricto,* known as hypervirulent *K. pneumoniae* (hvKp), cause severe infections in healthy members of the community. There is a paucity of research into rates of hvKp in the United States, the distribution of *K. pneumoniae* complex members in clinical specimens, and the pathogenic potential of complex members other than *K. pneumoniae sensu stricto.* We assessed the rates of hvKp and *K. pneumoniae* complex members in our 886 bed tertiary care facility in Rochester, New York. We performed whole genome sequencing on all inpatient, sterile-source isolates identified by routine culture work-up as *K. pneumoniae* from October 2018 – October 2019 (n=35). We additionally sequenced all *K. pneumoniae* liver isolates available in our collection, spanning 2017-2019 (n=18). In the yearlong collection of *K. pneumoniae* complex isolates, we found that 92.4% were *K. pneumoniae sensu stricto* (n=32), 5.7% were *K. quasipneumoniae* (n=2), and 2.9% were *K. variicola* (n=1). Five hvKp isolates were found, representing 5.7% (n=2) of *K. pneumoniae sensu stricto* isolates in the yearlong collection and 27.7% (n=5) of the liver isolate collection. All hvKp isolates were from liver; 60% were not associated with prior international residence.

## Background

*Klebsiella pneumoniae* is a major human pathogen that causes a wide variety of nosocomial and community acquired infections. It is the second most common cause of gram negative bacteremia and a leading cause of both urinary tract infections and ventilator-associated pneumonia [1–4]. Furthermore, infections caused by this organism can be particularly lethal; mortality rates of up to 50% have been reported [5–7]. While *K. pneumoniae* is notorious for being one of the ESKAPE pathogens, due to frequent multi-drug resistance and association with the production of KPC carbapenemases [8], there is also emerging awareness of other aspects of this pathogen, both in terms of phylogeny and virulence.

*K. pneumoniae* was recently determined to be a complex of closely related species. In addition to *K. pneumoniae sensu stricto,* the *K. pneumoniae* complex is composed of *K. variicola, K. quasipneumoniae, K. quasivariicola,* and *K. africana* [9–12]. Standard laboratory methods are generally unable to differentiate these species, though the most recent Bruker and Vitek2 MALDI databases allow *“Klebsiella pneumoniae”* and *“Klebsiella variicola”* to be distinguished [13–15]. Understanding of the epidemiology and clinical significance of individual members of the *K. pneumoniae* complex is still emerging. Systematic studies of *K. pneumoniae* complex clinical isolates have revealed that 2.8 – 24.5% of clinical isolates initially identified as *K. pneumoniae* are actually *K. variicola* upon sequencing analysis and 3 – 12.8% are *K. quasipneumoniae,* with considerable variance across studies [16–23]. Rates of antimicrobial resistance and evaluations of clinical severity of *K. variicola* and *K. quasipneumoniae* infections in these studies have similarly varied. Notably, *K. variicola* has been implicated in nosocomial outbreaks among neonates [24, 25]. There is very little known about the clinical significance of *K. quasivariicola* and *K. africana*, aside from three case reports of *K. quasivariicola* infections [26–28].

In addition to the recent discovery that *K. pneumoniae* represents a complex of closely related species, since the 1980’s hypervirulent *K. pneumoniae* (hvKp) strains have come to attention as a concerning emerging infection, predominantly in east Asia. These strains of *K. pneumoniae* are notable for causing devastating invasive infections in healthy members of the community [29]. Since its initial characterization, hvKp has become a leading cause of community acquired liver abscess, gram negative pneumonia and bacterial meningitis in Taiwan and is an emerging cause of necrotizing fasciitis [30–39]. HvKp has also been identified worldwide, [40] though there have been few systematic studies of hvKp in the United States [41–43].

Strains which cause hvKp are genetically characterized by the carriage of *rmpA* and/or *rmpA2,* regulators of capsule gene expression, possession of capsule type K1 or K2, and an expanded complement of siderophores, particularly aerobactin *(iuc)* [44–48]. Many of these virulence factors, including *rmpA*, *rmpA2*, and aerobactin, are carried on non-conjugative plasmids, the prototype of which is pLVPK [49, 50]. Molecular definitions of hvKp have varied over time, though recent studies have identified the simultaneous presence of the aerobactin gene cluster (*iuc*) and *rmpA* or *rmpA2* as a sensitive and specific marker for hvKp [43, 48]. It is generally believed that hvKp strains are *K. pneumoniae sensu stricto,* often of a single MLST, ST23. However, case reports of *K. variicola* and *K. quasipneumoniae* possessing genetic markers of hypervirulence do exist [17, 51–55].

In this study, we sequenced all tissue invasive isolates (n=35) identified as *“Klebsiella pneumonia”* by standard laboratory methods at our 886 bed tertiary care facility in Rochester, New York between October 2018 and October 2019 with the goal of determining the relative rates of isolation of *K. pneumoniae* complex members as well as their sequence types, antimicrobial resistance, and virulence factors associated with hvKp. To increase detection of hvKp, we focused further on liver isolates and sequenced all qualifying *K. pneumoniae* available in our strain collection going back to 1/2017 (n=18).

## Results

A total of 36 isolates were identified as *Klebsiella pneumoniae* from sterile source specimens from adult inpatients at Strong Memorial Hospital during the October 2018 – October 2019 time period. Only the first isolate per patient was included in this count. Of these, 35 isolates had been frozen and were available for sequencing. Sequencing, genome assembly, and annotation were successful carried out for all 35 isolates available. Specimens were predominantly from male patients (73%) with a mean age of 65 years (range: 40 – 85 years) (Table 1). Specimens were from sterile body fluids (n=15), liver (n=6), bone (n=3), kidney (n=3), other sterile tissues (n=3), joint or joint fluid (n=2), and CSF (n=1). 57% of the *K. pneumoniae* isolates were part of a polymicrobial infection, having been isolated with at least one other organism, 12% were phenotypic extended spectrum beta-lactamase (ESBL) producers, and none were carbapenem resistant (Table 1). An additional 12 isolates identified as *Klebsiella pneumoniae* isolated from liver were available in freezer collections dating back to 2017. Sequencing, genome assembly, and annotation were successful for all 12 of these isolates.

**Table 1.** Summary of Patient and Infection Characteristics for the October 2018 – October 2019 Collection

| Characteristic | Result |
| --- | --- |
| Patient Sex | 37% female (n=13) |
| Mean patient age | 65 years (range: 40 - 85 years) |
| Source | Sterile body fluid (n=15)<br>Liver (n=6)<br>Bone (n=3)<br>Kidney (n=3)<br>Other sterile tissue (n=3)<br>Joint/joint fluid (n=2)<br>CSF shunt (n=1) |
| Polymicrobial specimen | 57% (n=20) |
| Phenotypic ESBL | 12% (n=4, 34 tested) |
| Phenotypic CRE | 0% |

### Resolution of *K. pneumoniae* complex

Thirty-two isolates were found to cluster with *K. pneumoniae sensu stricto* (84.1% – 90.2% identity to reference) (Figure 1, Dataset S1). URMC-553 was found to cluster with *K. variicola* (92.2% identity to reference). URMC-555 and URMC-560 were found to cluster with *K. quasipneumoniae* (87.5 – 88.6%, identity to reference) (Figure 1, Dataset S1). These results correspond to a yearly isolation rate among tissue invasive isolates of 92.4% *K. pneumoniae sensu stricto* (n=32), 5.7% *K. quasipneumoniae* (n=2), and 2.9% *K. variicola* (n=1). Neither *K. quasivariicola* nor *K. africana* were identified among the strain set.

**Figure 1.**
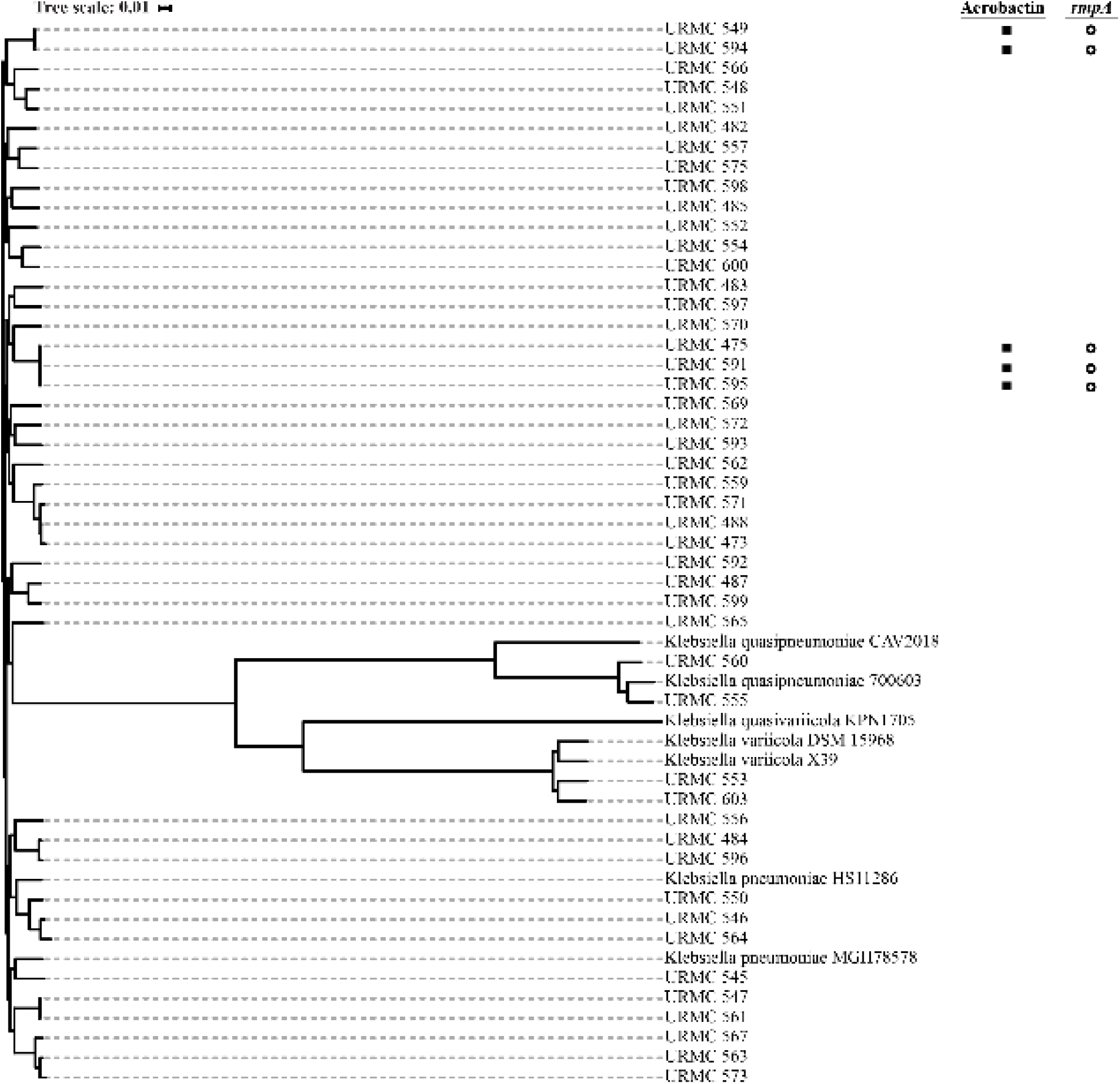
Phylogeny of *K. pneumoniae* isolates.

### *K. variicola* and *K. quasipneumoniae*

URMC-553 (*K. variicola)* was isolated in May of 2019 from a polymicrobial infection of joint fluid. It typed as ST729 with four variant loci according to the *K. pneumoniae* MLST scheme. Although a *K. variicola* specific MLST scheme has been developed, it remains standard practice to use *K. pneumoniae sensu stricto* MLST typing for all members of the *K. pneumoniae* complex [56]. URMC-553 carried the *bla*LEN-24 beta-lactamase, but had no other antimicrobial resistance genes identified (Dataset S2). Phenotypically, it was resistant to ampicillin and susceptible to all other antibiotics tested (Dataset S3).

The *K. quasipneumoniae* isolates, URMC-555 and URMC-560, were isolated in June 2019 from a polymicrobial liver abscess and October 2018 from a monomicrobial liver abscess, respectively. URMC-555 typed as ST414 in the *K. pneumoniae* MLST scheme. Phenotypically, it was resistant to ampicillin, ciprofloxacin, ceftriaxone, cefazolin, moxifloxacin, and trimethoprim-sulfamethoxazole, and intermediate to ampicillin/sulbactam and tobramycin (Dataset S3). An assessment of antibiotic resistance genes (ARGs) revealed genes encoding resistance to fluoroquinolones (*qnrS1*), rifampin (*arr-3*), sulfonamides *(sulI* and *sulII),* and trimethoprim *(dfrA27),* as well as the chromosomal beta-lactamase *bla*_OKP-B-8_ and the extended spectrum beta-lactamase *bla*_CRX-M-15_<. URMC-560 typed as ST1584. It carried the *sulI* gene encoding resistance to sulfonamides and the chromosomal beta-lactamase *bla*_OKP-B-10_. No other ARGs were detected. It was resistant to ampicillin and susceptible to all other antibiotics tested. (Dataset S3).

URMC-553, URMC-555 and URMC-560 were not found to carry genes associated with hypervirulence: genotypic hypermucoviscosity *(rmpA)* and the siderophore gene clusters for yersiniabactin *(ybt),* aerobactin (*iuc*), colibactin (*clb*), and salmochelin *(iro)* (Dataset S2).

### K. pneumoniae sensu stricto

The population structure of *K. pneumoniae sensu stricto* from the 2018 – 2019 sterile site collection was heterogeneous, separated by more than 200 SNPs across 26 STs (Dataset S2, Dataset S4).

Among isolates of *K. pneumoniae sensu stricto* on which antimicrobial susceptibility testing was performed, none were resistant to carbapenems. Four strains exhibited phenotypic ESBL activity (2.9%; URMC-488, URMC-561, URMC-547, and URMC-545) all of which carried *bla*_CTX-M-15_, except for URMC-488 in which a known ESBL gene was not detected. Two of these isolates, URMC-547 and URMC-561 were ST307, an international *bla*_CTX-M-15_ associated clone [57].

Given the low number of liver isolates in our initial collection, we expanded our criteria to include all liver isolates from 2017 – 2019 identified by conventional laboratory methods as *K. pneumoniae.* We found 12, all of which were available for sequencing. All 12 additional liver isolates were identified as *K. pneumoniae sensu stricto* (n=11) or *K. variicola* (n=1) by phylogeny (Dataset S1). Additional characteristics of these isolates can be found in Tables 2, S2 and S3.

### Hypervirulent *Klebsiella pneumoniae*

Two isolates from the initial 2018 – 2019 collection, URMC-475 and URMC-549, were determined to be hypervirulent by the presence of *rmpA/rmpA2* and the aerobactin gene cluster (*iuc*), for an overall yearly isolation rate of 5.3% (n= 2 of 35) (Table 3, Dataset S2). Both of these isolates were isolated from liver abscesses. Of the 12 additional liver isolates sequenced from 2017 – 2019, three, URMC-591, URMC-594, and URMC-595 were hypervirulent on the basis of the presence of *rmpA/rmpA2* and the aerobactin gene cluster, yielding a total of five hypervirulent isolates identified in this study. All five were K. *pneumoniae sensu stricto* (Table 4) and 4/5 were susceptible to all antibiotics tested except ampicillin (Dataset S3). URMC-595 was unique in also being resistant to nitrofurantoin (Dataset S3). All five isolates grew from pure or nearly pure cultures, had hypervirulence associated KL1 (URMC-475, URMC-591, URMC-595) or KL2 (URMC-549 and URMC-594) associated capsule types, and four (URMC-475, URMC-549, URM-591, and URMC-595) had hypervirulence associated ST types, with three (URMC-475, URMC-591, and URMC-595) being the epidemic ST23 (Table 4) [58]. Additionally, all five isolates carried sequences with homology to the full length of the pLVPK virulence plasmid.

**Table 2.** Summary of Patient and Infection Characteristics for Liver Isolate Collection (2017-2019)

| Characteristic | Result |
| --- | --- |
| Patient Sex | 16.7% Female (n=3) |
| Mean patient age | 62 years (range: 14 - 83) |
| Patient status | Inpatient: 18<br>Outpatient: 0 |
| Polymicrobial specimen | 66.7% (n=12) |
| Phenotypic ESBL | 5.6% (n=1) |
| Phenotypic CRE | 0% (n=0) |
| Species | *K. pneumoniae sensu stricto*: 83.3% (n=15)<br>*K. variicola*: 5.6% (n=1)<br>*K. quasipneumoniae*: 11.1% (n=2) |
| ST type (for ST types with n >1) | ST23: 16.7% (n=3) |

**Table 3.** Virulence Factors 2018-2019 Collection.

| **Virulence Factor** | **Percent of Isolates** |
| --- | --- |
| Genotypic hypermucoviscosity (*rmpA*) | 5.7 (n=2) |
| Yersiniabactin | 20 (n=7) |
| Aerobactin | 5.7 (n=2) |
| Colibactin | 2.9 (n=1) |
| Salmochelin | 5.7 (n=2) |
| Capsule type K1 | 2.9 (n=1) |
| Capsule type K2 | 8.6 (n=3) |

None of the hvKp patients experienced septic metastases, but three, URMC-549, URMC-591, and URMC-594, experienced treatment failures requiring readmission to the hospital. All five patients presented from the community. Two patients, URMC-591 and URMC-594, had anatomy that significantly predisposed them to liver infections (Table 4). URMC-549 and URMC-595 had a documented history of residence in the Asian Pacific Rim, where hvKp is endemic [40].

**Table 4.** Characteristics of Hypervirulent K. pneumoniae Isolates

| URMC ID | URMC-475 | URMC-549 | URMC-591 | URMC-594 | URMC-595 |
| --- | --- | --- | --- | --- | --- |
| Age | 48 | 83 | 74 | 24 | 66 |
| Sex | Male | Male | Male | Male | Male |
| Date of collection | 08/2019 | 03/2019 | 06/2018 | 06/2018 | 07/2018 |
| Polymicrobial | no | no | 1 colony *S. epidermidis* | no | no |
| Antimicrobial Resistance | no | no | no | no | nitrofurantoin |
| ST type | 23 | 373 | 23 | 86 single loci variant | 23 |
| Capsule type | KL1 | KL2 | KL1 | KL2 | KL1 |
| Siderophores | aerobactin<br>salmochelin | aerobactin<br>salmochelin,<br>yersiniabactin | aerobactin<br>salmochelin,<br>yersiniabactin, colibactin | aerobactin<br>salmochelin | aerobactin<br>salmochelin,<br>yersiniabactin, colibactin |
| Exposure to the Asian Pacific Rim | No evidence found | yes | No evidence found | No evidence found | yes |
| Predisposition | Impaired glucose tolerance | Chronic kidney disease | Whipple procedure | Hepatoporto enterostomy | None |

| Complications | None | 1x treatment<br>failure | 3x treatment<br>failure,<br>requiring<br>implant<br>removal | Multiple<br>treatment<br>failures, loss<br>of liver<br>function | None |
| --- | --- | --- | --- | --- | --- |

## Discussion

In our collection of sterile site specimens from between October 2018 and October 2019, we observed an isolation rate of 92.4% for *K. pneumoniae sensu stricto* (n=32), 5.7% for *K. quasipneumoniae* (n=2), and 2.9% for *K. variicola* (n=1). Our rates of *K. quasipneumoniae* are comparable to the 3 – 12.8% isolation rates previously reported from studies of blood, CSF, and respiratory specimens [16–23]. Our observed rates of *K. variicola* are low, but within the range (2.8 – 24.5%) that has been previously reported [16–23]. There was no evidence of clonal spread within our collection, with all isolates differing by more than 200 SNPs (Dataset S4) [59].

Our initial sample set suggested that *K. quasipneumoniae* was enriched among our *K. pneumoniae* complex liver isolates (33%), though the overall number was very small (n=2). Most previous reports of clinical *K. quasipneumoniae* isolates were isolated from bloodstream infections, with a lesser proportion isolated from urine, respiratory secretions, wound infections, and CSF. Both of the patients from whom our initial *K. quasipneumoniae* isolates were cultured had corresponding positive blood cultures identified by conventional work up as *K. pneumoniae*. As our conventional culture work up cannot differentiae *K. pneumoniae sensus stricto* from *K. quasipneumoniae*, it is likely that these positive blood cultures were also *K. quasipneumoniae*. This raised the possibility that tissue invasive *K. quasipneumoniae* may have a tropism for the liver, though neither of these isolates harbored virulence factors associated with hvKp (Dataset S2). An expanded analysis of *K. pneumoniae* complex liver isolates (2017 – 2019) in our laboratory failed to reveal additional *K. quasipneumoniae* isolates. Furthermore, a comprehensive review of English language literature revealed only one previous report of *K. quasipneumoniae* isolated from liver, so our exclusive isolation of *K. quasipneumoniae* from liver specimens remains somewhat unexpected [54].

We observed an isolation rate of 5.7% (n=2) for hvKp in our collection of sterile site specimens from Strong Memorial Hospital from between October 2018 and October 2019. This rate is consistent with the limited previous studies of the rate of hvKp in North American hospitals, with a rate of 8.2% found in Calgary, Alberta, Canada, rates of 7.9% and 3.9% in different hospitals in Houston, Texas, USA, and a rate of 3.7% in a New York City, New York, USA hospital [42, 43, 60]. Two of these studies, Periano et al. and Chou et al., collected only blood isolates, while Parrott et al. collected isolates from a variety of clinical specimens, predominantly blood.

In our expanded analysis of *K. pneumoniae* complex liver isolates in our laboratory, we identified three further hvKp strains, for a total of five. The microbiological characteristics of URMC-475, URMC-549, URMC-591, URMC-594, and URMC-595 were consistent with typical hvKp strains: encoding *rmpA/rmpA2,* aerobactin, and salmochelin, generally susceptible to all antibiotics except ampicillin, capsule type KL1 or KL2, and recovered from a pure or essentially pure culture. All of the hvKP isolates were *K. pneumoniae sensu stricto.*

HvKp infections classically present as liver abscess complicated by hematogenous seeding of other organs in up to a quarter of cases and development of severe endophthamitis leading to blindness in up to 11% of cases [37, 61]. Mortality in these patients is approximately 5-10% [61]. Meningitis, necrotizing pneumonia, and necrotizing fasciitis are also common manifestations of hvKp [40, 62]. Three of our hvKp patients fit the clinical definition of *K. pneumoniae* primary liver abscess syndrome (KLA): *K. pneumoniae* liver abscess occurring in the absence of predisposing intraabdominal factors (hepatobiliary disease, colorectal disease, history of intraabdominal surgery) [63]. Patient URMC-591 had a history of the Whipple procedure and had a biliary stent in place. Additionally, isolate URMC-591 was recovered in mixed culture with *Staphylococcus epidermidis.* Patient URMC-594 had a history of biliary atresia treated by hepatoportoenterostomy in infancy. Although these two cases do not fit the clinical definition of KLA, these isolates should not be discounted as hvKp due to their isolation from patients with predisposing conditions. None of our patients experienced septic metastases. This is not unusual among reports of hvKp infections and KLA in North America. Even in Taiwan, where septic metastases are frequently reported, they occur in only a minority of patients with hvKP infections, and thus the lack of them in this small case series should not be considered unusual [37, 62].

Management of hvKp infections is generally the same as management of classical *K. pneumoniae* infections, though special considerations include the need to use large bore catheters for draining abscesses (as the hypermucoviscous nature of hvKp can cause standard catheters to clog) and the need for enhanced imaging studies to detect septic metastases. Antimicrobial resistance among hvKp strains generally remains rare. However, in some studies rates of extended spectrum beta-lactamase producing (ESBL) hvKp have been as high as 13% and case reports of carbapenemase producing hvKp have emerged [39, 64–68]. It is worth noting that three of the five patients with hvKp experienced at least one treatment failure requiring repeat hospitalization, underscoring the need for specialized management of hvKp infections.

HvKp accounted for 30.0% of all *K. pneumoniae sensu stricto* isolated from liver in our lab between 2017 and 2019 (n= 5 of 15), demonstrating that it accounts for a substantial minority of our *K. pneumoniae sensu stricto* liver isolates. Of these isolates, both URMC-549 and URMC-595 came from patients with clear evidence of residence in or travel to the Asian Pacific Rim, where hvKp is endemic [40]. Given that asymptomatic intestinal carriage of hvKp occurs in healthy individuals in endemic areas, it seems likely that these two patients acquired their hvKp strain during the period of their international residence [43]. While it is impossible to rule out that patients URMC-475, URMC-591 and URMC-594 may have a relevant travel history that was not recorded in their medical records, the lack of such history in 60% of the patients with hvKp suggests that hvKp is endemically circulating in the URMC catchment area.

In summary, the rates of *K. pneumoniae sensu stricto*, *K. quasipneumoniae*, and *K. variicola* observed in this study are consistent with those observed internationally and the rates of hvKp are consistent with those seen at other hospitals in North America.

## Methods

### Ethics statement

Testing of the *Klebsiella* isolates and limited collection of patient data were approved by the Institutional Review Board of the University of Rochester Medical Center (URMC). For all isolates, patient data on age, sex, hospital site admitted to and inpatient/outpatient status were collected from the laboratory information system. For patients with hvKp, *K. quasipneumoniae*, or *K. variicola* isolates, a full review of the electronic medical record was undertaken to collect information on preexisting conditions, clinical outcomes and complications, and documented history of foreign travel or residence.

### Study site and samples

The URMC is an 886 bed tertiary care facility located in Western New York which services a large catchment area which includes the Rochester, New York metropolitan area and the surrounding rural counties. The URMC Central Laboratories additionally provides reference microbiological services to a larger catchment area which includes several small community hospitals and encompasses much of Western New York.

Two sets of criteria were used for the inclusion of isolates in this study. For the first set of criteria, all isolates identified by routine culture work up as *K. pneumoniae* that were isolated from sterile site specimens collected from inpatients at Strong Memorial Hospital between October 2018 and October 2019 (n=35) were selected for inclusion. For the second, all isolates identified by routine culture work up as *K. pneumoniae* isolated from liver specimens available at the URMC Central Laboratory and not already included in the October 2019 – October 2019 collection were selected for inclusion in the study (n=12).

Prior to July 2019, isolate identification was accomplished using the Biomerieux Vitek MS Matrix Assisted Laser Desorption Ionization (MALDI) Time-of-Flight system. Following July 2019, identification was accomplished using the Bruker MALDI Biotyper System. For isolates isolated prior to July 2019, antibiotic susceptibility test was accomplished using the Biomerieux Vitek 2 Gram negative antimicrobial susceptibility testing (AST) card. For isolates isolated after July 2019, antibiotic susceptibility testing was accomplished using the BD Phoenix NMIC-300 panel.

### DNA extractions and whole genome sequencing

Isolates were subcultured from frozen stocks onto BD BBL Trypticase Soy Agar with 5% Sheep Blood plates. Overnight growth was lysed with lysozyme in Roche MagNA Pure Bacteria Lysis Buffer for 1 hour. Total nucleic acid was extracted from the lysate using the Roche MagNA Pure Compact Nucleic Acid Isolation Kit I and the MagNA Pure Compact instrument (Roche, Indianapolis, Indiana). Dual-indexed sequencing libraries were prepared from the lysates using Nextera XT DNA Library Preparation Kit (Illumina FC-131-1096), and sequenced on an Illumina Miseq benchtop sequencer (Illumina, San Diego, CA) using Illumina MiSeq reagent kit V3, 600 cycle.

### Genomic Analysis

Genome assembly: Genome assembly and analysis was performed using the URMC Bacterial Genomic Analysis Pipeline version 3.0.3 (isolates URMC-473, −475, −482, −483, −484, −485, −487, −488, −545, −546, −547, −548, −549, −550, −551, −552, −553, −554, −555, −556, −557, – 559, −560, −561, −562, −563, −564, −565, −566, −567) or the URMC Bacterial Genomic Analysis Pipeline version 3.1.1 (isolates URMC −569, −570, −571, −572, −573, −575, −591, −592, −593, −594, −595, −596, −597, −598, −599, −600, −603), a previously published in-house biofinformatics pipeline written in Python, sqlite3, BASH, JavaScript, D3, JQuery, HTML, and Bootstrap and run on high-performance computer cluster at the Center for Integrated Research Computing at the University of Rochester [69, 70]. Updates between version 3.0.3 and 3.1.1 addressed ease of use and did not impact pipeline functionality. Details of each step in the pipeline are provided below.

Read quality control: On average, 20 human reads were found contaminating raw reads which were removed using Bowtie2 [71]. Trimmomatic *v*0.36 was used to ensure high quality reads [72]. Trimmomatic default conditions were set to (LEADING:3 TRAILING:3 SLIDINGWINDOW:1:20 MINLEN:50). Quality scores across all bases were checked with FastQC v0.11.5 [73]. On average there were 701,079 forward and reverse reads which passed quality control with an average length of 254 for the forward reads and 224 for the reverse reads.

Sequence assembly and mapping: De novo assemblies and reference alignments were both built from reads which passed the above quality control metrics. Genome assemblies with an average N50 of 243,452.9 were built with using SPAdes v3.11.1 using the --careful setting [74]. Quality of genome assembly and genome fraction for each reference was assessed by Quast v4.5 [75]. References were aligned against the *K. pneumoniae sensu stricto* strain MGH78758 (NCBI: NC_009648.1) using bowtie2 v2.2.9 [71]. Alignment resulted in an average coverage of 86X and an average depth of 55X (excluding regions below 12X). Annotation was accomplished using Prokka version 1.12 [76].

### SNP Calling and Phylogenetic Analysis

A modified CFSAN SNP Pipeline v.1.0.0 was used for reference-based SNP Calling and Phylogenetic Analysis [77]. CFSAN SNP Pipeline call_sites uses samtools v1.5 and varscan (min-var-freq:0.90, min-coverage:12, min-avg-qual:20) to find the sites having high-confidence SNPs between the reference and each of the mapped reads of individual samples [78, 79]. Abnormally dense SNPs are removed making a preserved_vcf file using the CFSAN SNP Pipeline filter_regions (window_size: (default 50), max_num_snps: (default3)). All high-confidence SNP positions across the samples are combined into a single SNP list using CFSAN SNP pipeline merge_sites. The default run conditions were set to stop the practice of removing SNPs inside of phages, mobile elements, and transposons based on a RAST vClassicRAST annotated gff file [80]. A concatenated SNP fasta file for each sample was produced using CFSAN SNP pipeline call_consensus (minConsFreq: 0.9). Where a consensus could not be found for a sample, a dash was inserted into the fasta sequence. Default run conditions were set to stop the practice of finding a common backbone of concatenated SNPs from each FASTA sequence producing a multi-FASTA file without dash loci. A maximum likelihood tree was produced from the multi-FASTA file with FastTree [81]. FastTree v2.1.10 produces a newick formatted text file which is visualized using FigTree [82]. The CFSAN SNP Pipeline uses the multi-FASTA file to produce SNP distance matix. A matrix of samples names as columns vs SNP genome location, gene name based on RAST annotation, gene length, gene location as rows was produced. Values in the matrix can either be none or mutation change based on the comparison of each sample with the reference. A web application was used to visualize the coverage and SNP locations throughout the genome to ensure consistent coverage and no SNP clustering. The online tool, the Interactive Tree of Life (iTOL), was used to visualize the maximum likelihood tree after data had been deidentified [83]. Phylogeny was determined on the basis of phylogenetic clustering using parsnp with filtering of SNPs located in PhiPack identified regions of recombination, a curated genome directory, and 20 threads [84]. Species assignment was based on the following reference genomes: *K. pneumoniae sensu stricto* MGH78758 (NCBI: NC_009648.1), *K. pneumoniae sensus stricto* HS11288 (NCBI: NC_016845.1), *K. quasipneumoniae* CAV2018 (NCBI: NZ_CP029432.1), *K. quasipneumoniae spp. similipneumoniae* 700603 (NCBI: NZ_CP014696.2), *K. quasivariicola* KPN1705 (NCBI: NZ_CP022823.1), *K. variicola* DSM15968 (NCBI: NZ_CP010523.2), and *K. variicola* X39 (NCBI: NZ_CP018307.1). No reference genome was available for *K. africana*, so it was omitted.

### Prediction of virulence and antimicrobial resistance genes

The bioinformatics pipeline Kleborate version 0.2.0 (https://github.com/katholt/Kleborate) was used with default parameters and --kaptive and --resistance enabled to determine sequencing type, determine K locus type, identify virulence factors, and identify antibiotic resistance genes [85–88]. Isolates found to simultaneously contain the aerobactin operon and *rmpA* or *rmpA2* were classified as hypervirulent. Presence of the pLVPK virulence plasmid (NC_005249.1, Klebsiella pneumoniae CG43 plasmid pLVPK, complete sequence) in each isolate was assessed using nucleotide BLAST (max target sequences set to 100, short queries enabled, expect threshold set at 10, maximum matched in a query range set at 0, match/mismatch scores set at 1 at −2, respectively, gap costs set to linear, filter low complexity regions enabled, masking set to lookup table only) to query the full plasmids sequence (NC_005249.1) against the sequences of each isolate [89].

## Data availability

All genomes were deposited in the NCBI database under Bioproject PRJNA673499. Reference numbers for each strain can be found in Dataset S5.

## Acknowledgements

This work was funded by the University of Rochester Department of Pathology and Laboratory Medicine. The authors would like to thank Zachary Pearson for assistance with pulling isolates.

